# Identification of 17 novel epigenetic biomarkers associated with anxiety disorders using differential methylation analysis followed by machine learning-based validation

**DOI:** 10.1101/2024.05.23.595430

**Authors:** Yoonsung Kwon, Asta Blazyte, Yeonsu Jeon, Yeo Jin Kim, Kyungwhan An, Sungwon Jeon, Hyojung Ryu, Dong-Hyun Shin, Jihye Ahn, Hyojin Um, Younghui Kang, Hyebin Bak, Byoung-Chul Kim, Semin Lee, Hyung-Tae Jung, Eun-Seok Shin, Jong Bhak

## Abstract

**Background:** The changes in DNA methylation patterns may reflect both physical and mental well-being, the latter being a relatively unexplored avenue in terms of clinical utility for psychiatric disorders. In this study, our objective was to identify the methylation-based biomarkers for anxiety disorders and subsequently validate their reliability.

**Methods:** A comparative differential methylation analysis was performed on whole blood samples from 94 anxiety disorder patients and 296 control samples using targeted bisulfite sequencing. Subsequent validation of identified biomarkers employed an artificial intelligence-based risk prediction models: a linear calculation-based methylation risk score model and two tree-based machine learning models: Random Forest and XGBoost.

**Results:** 17 novel epigenetic methylation biomarkers were identified to be associated with anxiety disorders. These biomarkers were predominantly localized near CpG islands, and they were associated with two distinct biological processes: 1) cell apoptosis and mitochondrial dysfunction and 2) the regulation of neurosignaling. We further developed a robust diagnostic risk prediction system to classify anxiety disorders from healthy controls using the 17 biomarkers. Machine learning validation confirmed the robustness of our biomarker set, with XGBoost as the best-performing algorithm, an area under the curve of 0.876.

**Conclusion:** Our findings support the potential of blood liquid biopsy in enhancing the clinical utility of anxiety disorder diagnostics. This unique set of epigenetic biomarkers holds the potential for early diagnosis, prediction of treatment efficacy, continuous monitoring, health screening, and the delivery of personalized therapeutic interventions for individuals affected by anxiety disorders.

## Background

Anxiety disorders, with a global prevalence of approximately 4%, have a higher prevalence compared to other mental disorders.[1] Despite their high prevalence, anxiety disorders are challenging in many regards: diagnosis, primary care, treatment effectiveness monitoring, and relapse prevention.[2] These diverse challenges stem from the difficulty of accurately measuring the state of mental health in individuals, given that mental disorders arise from the complex interplay of genetic and environmental factors, making it challenging with conventional techniques.[3] A twin study suggested that environmental factors may account for up to 12% of clinical variance in anxiety disorders.[3] Moreover, traditional diagnostic methods relying on questionnaires may exhibit variability contingent upon the subject’s health status or mood during assessment, underscoring the imperative for complementary approaches.[4, 5] These insights highlight the need for novel approaches that could directly measure these complicated effects from molecular materials derived from the patient, circumventing biases that standard clinical diagnostic questionnaires and interviews are prone to. Recent genomic studies for anxiety disorders, although not reaching any definitive conclusions yet, have yielded a range of diverse biomarkers from various tissues. For example, Edelmann *et al*., investigated the blood transcriptome and identified 13 significantly differentially expressed genes (DEGs) between individuals diagnosed with social anxiety disorder and healthy controls.[6] Levey *et al*., reported five blood-derived SNPs associated with Generalized Anxiety Disorder 2- item (GAD-2) scale for European Americans and one SNP in African Americans, respectively,[7] while Su *et al*., identified 26 genes associated with anxiety through a transcriptome-wide association study (TWAS) utilizing multiple brain tissues.[8] Two blood-derived differentially methylated regions (DMRs) had been reported in association with social anxiety disorder by Wiegand et al. in 2020.[9] Furthermore, in 2023, an epigenome-wide associdation study (EWAS) was conducted on the methylation patterns of anxiety disorder patients based on their blood cell types.[10] However, prior studies faced critical limitations in validating identified biomarkers, as well as lacked dedicated research exploring their application in diagnostic scenarios.

In order to attain a comprehensive understanding of anxiety disorders, which are influenced by a combination of genetic and environmental factors, the utilization of epigenetic data reflecting long-term environmental influences is critical.[11, 12] A particularly practical approach in this regard involves the assessment of DNA methylation changes in readily accessible tissues, such as blood, for diagnostic purposes. This approach facilitates the integration of epigenetic insights into clinical practice, thus offering valuable insights into the underlying molecular mechanisms and potential diagnostic and therapeutic strategies for anxiety disorders.

This study shows potential for employing peripheral blood and molecular diagnostics in diagnosing anxiety disorders. Bootstrapping techniques were employed to identify precise methylation biomarkers in 94 Korean patients diagnosed with anxiety disorders by physicians. Following this, we developed machine learning models to validate the diagnostic capabilities of these biomarkers, assessing their performance in classifying cases from controls in the validation set.

## Methods

### Study population

A total of 94 patients with anxiety disorder from the Ulsan Medical Center, Ulsan, Korea were enrolled in this study, and 296 healthy individuals from the Korean Genome Project (KGP) were selected as controls.[13, 14] The healthy control group was configured to match the sex and age distribution of the anxiety patient group (Table 1). All of our anxiety disorder patients received their diagnosis from a psychiatric specialist in a clinical setting. All samples used in our study are presumed to be ethnically Koreans. Information regarding the KGP data can be found on the Korean Genome Project webpage (http://koreangenome.org).

**Table 1.** Baseline characteristics in this study.

|  | Anxiety<br>disorder patient | Healthy control<br>(n=296) | *p*-value |
| --- | --- | --- | --- |

|  |  | (n=94) |  |  |
| --- | --- | --- | --- | --- |
| **Sex** | **Male** | 41 (43.62 %) | 134 (45.27 %) | 0.813 |
|  | **Female** | 53 (56.38 %) | 162 (54.73 %) |  |
| **Age** |  | 42.98 (15.64) | 40.40 (15.76) | 0.168 |
| **Type of anxiety disorder** | **Panic disorder** | 10 (10.64 %) | 0 | NA |
|  | **Social anxiety disorder** | 6 (6.38 %) | 0 |  |
|  | **Generalized anxiety disorder** | 78 (82.98 %) | 0 |  |

Continuous values are represented as mean (standard deviation). Proportions are shown as absolute number (percentage). The following statistical methods were applied: (Sex: Fisher's exact test, Age: Student's t-test, Type of anxiety disorder: Chi-square test). NA = Not Available;

### Study design

This study was performed to discover and validate differential methylation biomarkers between anxiety disorder patients and healthy controls, employing a machine learning-based approach (Fig. 1). The entire dataset was divided into training and validation sets, ensuring even distribution of sex, age, and types of anxiety disorders in each set (Table 1, Additional file 3: Table S1).

**Fig. 1.**
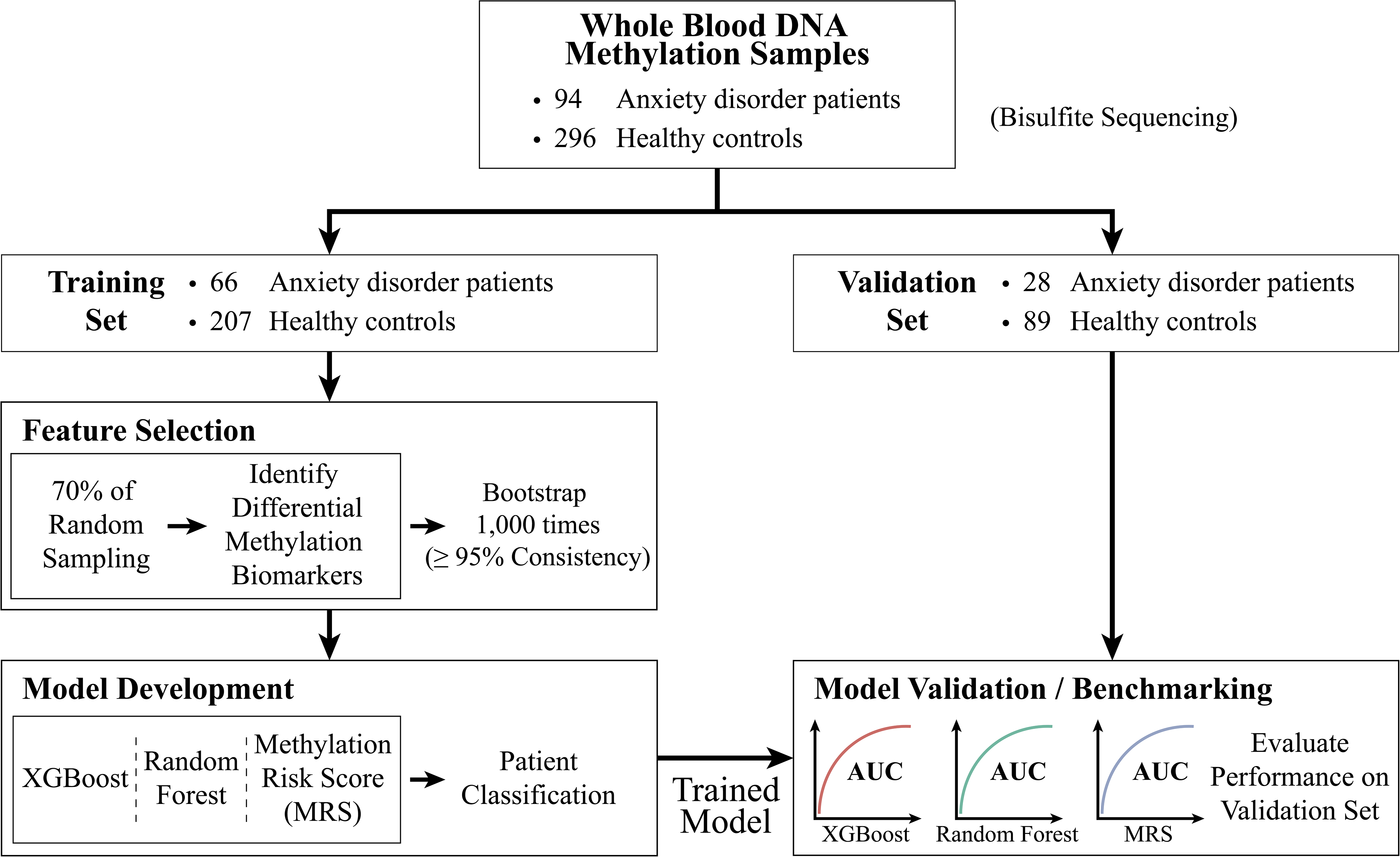
Study design for the discovery and validation of differentially methylated biomarkers for anxiety disorders. The flow chart illustrates the utilization of methylation data according to the flow of arrows. AUC = Area Under Curve;

### Targeted bisulfite sequencing and raw data processing

Blood samples were collected in EDTA tubes at the time of recruitment, then frozen, and genomic DNA was extracted using the DNeasy Blood and Tissue Kit from Qiagen (Qiagen, 69506), following the manufacturer’s protocol. The library preparation process followed the protocol of the SureSelectXT Methyl-Seq Target Enrichment System Kit (Agilent, G9651). Then, sequencing was performed using both the Illumina NovaSeq 6000 platform and the Hiseq 2500 platforms, generating paired-end reads of 150bp and 100bp, respectively. The adapter trimming process for the sequenced reads was performed using the fastp (version 0.23.1) with specific parameters (thread=16, trim_front1=15, trim_front2=15, trim_tail1=5, trim_tail2=5, length_required=0, cut_front=1, cut_tail=1, cut_right=0, cut_mean_quality=20, average_qual=20, n_base_limit=1, detect_adatper_for_pe).[15, 16] To analyze the results of bisulfite sequencing, we utilized Bismark (version 0.23.1) with Bowtie2 (version 2.5.1) to align the bisulfite-treated reads to the reference genome (hg38) without scaffolds.[17, 18] We applied the parameters “--L 20 -N 1 -p 10 -q –phred33_quals --dovetail” during the alignment process. Methylation calling for CpG sites was then performed with Bismark (version 0.23.1).[17]

### Identification of methylation biomarkers for anxiety disorders

We used the methylKit package (R version 4.1.3; methylKit version 1.20.0) to identify methylation biomarkers in individuals with anxiety disorders.[19, 20] For each CpG site, if the

sequencing depth was less than 10, it was treated as NA (not available). 862,525 CpG sites with values present in all samples were utilized for the analysis. In order to identify methylation biomarkers able to distinguish healthy controls from patients with anxiety disorders, a logistic regression was conducted using the methylKit package, incorporating sex and age as covariates. Furthermore, overdispersion correction was utilized to enhance the precision of the biomarkers, removing false positives.[21] Significant biomarkers were identified using a False Discovery Rate(FDR; Benjamini-Hochberg correction)-adjusted *p*-value cutoff of 0.05 and at least 10% of mean difference in methylation between patient and control groups in DNA methylation as the threshold.

### Establishing a Machine learning Model for Validating the Performance of Biomarkers

For the purpose of validating the performance of methylation risk scores or machine learning models, the entire dataset was partitioned into a training set and a validation set using a 7:3 ratio, facilitating the extraction of biomarkers. The training and validation sets were stratified throughout this procedure to ensure parity in sex and age distributions (Additional file 3: Table S1). Moreover, exclusive reliance on the training set was employed in the biomarker discovery process to mitigate overfitting concerns in feature selection. A 70% random sampling within the training set was conducted 1,000 times to mitigate potential bias due to the reduced training set size. Subsequently, we performed methylation biomarker discovery for 1,000 times using the previously outlined methods. Biomarkers that consistently appeared in 95% or more instances across the 1,000 independent analyses were considered robust and eligible for further analysis.

To optimize the biomarker set, we applied various thresholds for the reproducibility rate in the 1,000 biomarker discovery iterations (95%, 96%, 97%, 98%, 99%, and 100%) (Additional file 2: Fig. S1A, and S1B).

### Calculation of methylation risk score for anxiety disorders

To assess the performance of the biomarkers identified in the training set, we calculated the Methylation Risk Score (MRS) as an indicator for classifying anxiety disorder patients and healthy controls in the validation set. In the MRS calculation, logistic regression (1) was conducted on the training set to determine the weight(w^CpG^) of each i-th biomarker as follows:

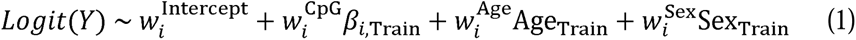

where Y corresponds to the condition (Healthy control : 0, Anxiety disorder patient : 1), Age and Sex the covariates of age and sex for each sample in the training set, ,B the methylation value of the i-th biomarker for the training set, w the coefficient for each variable.

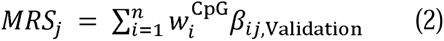

For each sample j, the MRS encompassing the entire set of n methylation biomarkers was computed by iteratively multiplying the coefficient and methylation value (,B_ij_) derived from the logistic regression model for each i-th CpG site and subsequently summing these products (2). The MRS formula was applied to the validation set for the validation of methylation biomarkers.

Subsequently, the performance of the MRS was evaluated by analyzing ROC (Receiver operating characteristic) curves.

### Application of machine learning models based on differential DNA methylation

The Random Forest model was trained using Sci-kit learn package (version 0.23.2), and the XGBoost model was trained using xgboost package (version 1.6.2). Hyperparameter tuning was conducted using GridSearchCV on the validation set (Additional file 4: Table S2, and Additional file 5: Table S3). Both models underwent a 3-fold cross-validation during the hyperparameter tuning process. In the hyperparameter optimization process, we first established the initial values for all parameters and subsequently conducted a more detailed optimization for each parameter individually. For the XGBoost model, the tuning sequence was performed in the following order: max_depth, min_child_weight, gamma, subsample, colsample_bytree, and n_estimators (Additional file 4: Table S2). In the Random Forest model, the tuning was executed in the following order: max_depth, min_samples_split, min_samples_leaf, and n_estimators (Additional file 5: Table S3). To compute the feature importance of each biomarker in the model, we calculated the SHapley Additive exPlanations (SHAP) using the shap package (version 0.41.0).[22] The entire procedure was carried out in a Python 3.7.16 environment.

### Annotation of methylation biomarkers

We annotated the differentially methylated sites (DMSs) using Annotatr (R version 4.1.3; Annotatr version 1.20.0)[23], utilizing the options "hg38_cpgs," "hg38_basicgenes," and "hg38_enhancers_fantom" for the annotation of each CpG site. If a particular CpG site was associated with the location of more than two genes, only those genes related to the promoter region were annotated. Utilizing the ClueGO plugin (version 2.5.9)[24] in Cytoscape (version 3.9.1), we conducted functional enrichment analysis in Gene Ontology (GO, http://geneontology.org/)[25, 26] and Kyoto Encyclopedia of Genes and Genomes (KEGG, https://www.genome.jp/kegg/).[27–29] The *p*-values from the enrichment test underwent correction using the Benjamini-Hochberg method to calculate the False Discovery Rate (FDR). We identified statistically significant entries based on the criterion of FDR < 0.05.

### Statistical analysis

All the statistical analyses were performed using the SciPy package (version 1.7.3) in Python (version 3.7.16). Fisher’s exact test assessed the difference in sex ratio between healthy control and anxiety disorder patients, while Student’s t-test examined the difference in age distribution between them. To verify the equivalent stratification of samples during the train-validation splitting, the Chi-square test checked the distribution of the type of anxiety disorder among anxiety disorder patients in the training and validation sets. Furthermore, Fisher’s exact test and Student’s t-test were applied to evaluate the distribution of each sex and age between training and validation sets. Additionally, the Mann-Whitney-Wilcoxon test was conducted to assess significant differences in methylation across specific types of anxiety disorder.

## Results

### Detecting differentially methylated sites and classification model construction

Employing bootstrapping on the training set to maximize the robustness, we identified 18 methylation biomarkers that showed significantly different methylation values with over 95% reproducibility out of 1,000 biomarker discoveries between 66 anxiety disorder patients and 207 healthy controls (Table 1, Fig. 1, Fig. 2A, and Additional file 2: Fig. S1A-C, Additional file 6: Table S4). The optimal marker set, however, consisted of 17 markers that exhibited a reproducibility of at least 97%, reflecting the consistency in marker discovery during bootstrapping (Fig. 2B, 2C, Additional file 2: Fig. S2A, and S2B). Of these, 16 were hypo- methylated biomarkers, and one (CpG site located in the intronic region of the *INPP5A* gene) was hyper-methylated. Even though these biomarkers were discovered in the training set, they were also significantly different between anxiety disorder patients and healthy controls in the validation set as well (17 markers *p*-value < 0.05 on both train and validation set) (Additional file 2: Fig. S1D-F).

**Fig. 2.**
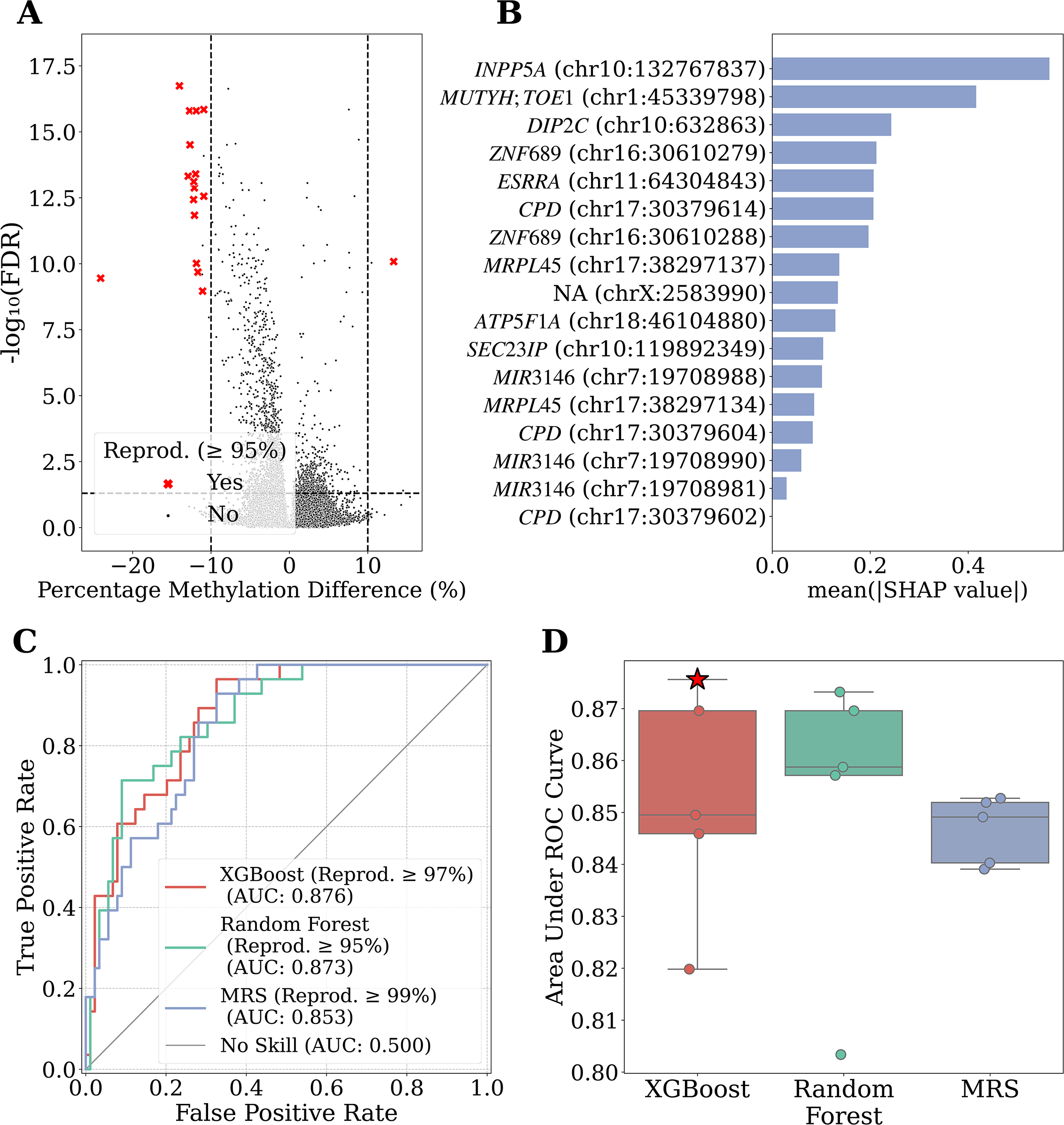
Discovery of Biomarkers Predictive of Anxiety Disorders and Construction of Classification Model. A) Volcano plot depicting biomarkers observed in the entire dataset. Biomarkers showing equal to or higher than 95% reproducibility through bootstrapping in the training set are highlighted in red. Reprod. = Reproducibility; B) The feature importance of each biomarker using the best performing model (XGBoost) on the consensus biomarker set with 97% of reproducibility. The X-axis represents the average impact on the model output magnitude, while the Y-axis depicts the positions of 17 biomarkers showing 97% or higher reproducibility and the associated gene names. If there are two or more genes associated, they are separated by a semicolon. SHAP = SHapley Additive exPlanations; C) Receiver Operating Characteristic (ROC) curves for each model (XGBoost, Random Forest, and MRS) using the biomarker set that demonstrated the highest performance. Reprod. = Reproducibility; D) Box plot illustrating the Area Under the ROC curve (AUROC) values for each model (XGBoost, Random Forest, and MRS). Each dot represents the performance of the model for each of the five reproducibility thresholds, and the star indicates the model with the highest performance among all conditions.

To evaluate the discriminatory power of 18 methylation biomarkers on 28 anxiety disorder patients and 89 healthy controls, three primary approaches were employed: a linear calculation-based MRS model and two tree-based machine learning models (Random Forest and XGBoost) (Fig. 1). Overall, all models demonstrated high performance with AUROC (Area under the receiver operating characteristic curve) above 0.8 (Fig. 2D). In the MRS model, the performance differences across biomarker sets (subsampled from the 18 identified putative biomarkers based on their reproducibility) were relatively consistent (Standard deviation of AUROC for MRS : 0.0058), while machine learning-based models exhibited more considerable variations (Standard deviation of AUROC for Random Forest : 0.0253, XGBoost : 0.0197) (Additional file 2: Fig. S2C). The average performance, ranked from highest to lowest based on AUROC, was observed as follows: Random Forest, XGBoost, and MRS (Mean AUROC of Random Forest : 0.8524, XGBoost : 0.8521, and MRS : 0.8466). The overall high performance of these models indicates the robustness of the markers identified in this study in classifying anxiety disorders, regardless of the method used.

This marker set was further optimized using the best performing model, XGBoost, with multiple reproducibility thresholds and yielded an even more robust 17 biomarker set. Subsequently, training the XGBoost model with 17 markers exhibiting a reproducibility of 97% or higher based on AUROC values yielded the highest performance (AUROC : 0.876, AUPRC : 0.654, Sensitivity : 96.429%, Specificity : 67.416%) (Additional file 2: Fig. S2A, and S2B). This model demonstrated the highest performance compared to previous models for classifying anxiety disorder patients based on electronic health check-up records (AUROC : 0.73, Sensitivity : 66%, Specificity : 70%)[30], brain gray matter volumes in adolescents (AUROC : 0.63)[31], or the DASS-21 (The Depression, Anxiety and Stress Scale - 21 Items) questionnaire (73.3% accuarcy on Naïve Bayes model).[32] In this final model, we calculated the feature importance of each biomarker using SHapley Additive exPlanation (SHAP) values, which revealed the only hyper- methylated CpG site, located in the intronic region of the *INPP5A* gene, had the highest importance (SHAP = 0.5658), followed by the hypo-methylated intergenic CpG site located in between of the *MUTYH* and *TOE1* genes (SHAP = 0.4163) (Fig. 2B).

### Four hypomethylation biomarkers show potential for distinguishing the type of anxiety disorder

Our anxiety disorder patient group is rather heterogeneous; comprised of individuals with three distinct diagnoses: panic disorder (PD), social anxiety disorder (SAD), and generalized anxiety disorder (GAD). Consequently, the 17 biomarkers previously validated for general use were subjected to test their specificity towards each type of disorder (Additional file 2: Fig. S4). All the biomarkers proved effective in distinguishing patients with GAD from healthy controls (All 17 markers *p*-value < 1×10^-8^) (Additional file 2: Fig. S4, and Additional file 7: Table S5). In contrast, for six CpG sites, the methylation values of patients with panic disorder or social anxiety disorder did not significantly differ from those of healthy controls (Additional file 2: Fig. S4G, S4H, S4I, S4J, S4N, S4Q, and Additional file 7: Table S5).

Among the 17 CpG sites, four CpG sites exhibited significant differences in the detailed classification of anxiety disorders (*p*-value < 0.05; Fig. 3, Additional file 2: Fig. S4, and Additional file 7: Table S5). Three of these biomarkers were located in the promoter region of the *CPD* gene on chromosome 17 (Fig. 3A-C). These biomarkers did not show significant differences between patients with panic disorder or social anxiety disorder and healthy controls; instead, they exhibited significant differences when compared to patients with generalized anxiety disorder (Fig. 3A-C). This pattern of methylation changes indicates specific alterations unique to GAD. This is influenced by the predominant representation of generalized anxiety disorder patients, overshadowing the relatively fewer cases of panic disorder and social anxiety disorder. Therefore, it is possible that the majority of DMSs followed the trend of generalized anxiety disorder.

**Fig. 3.**
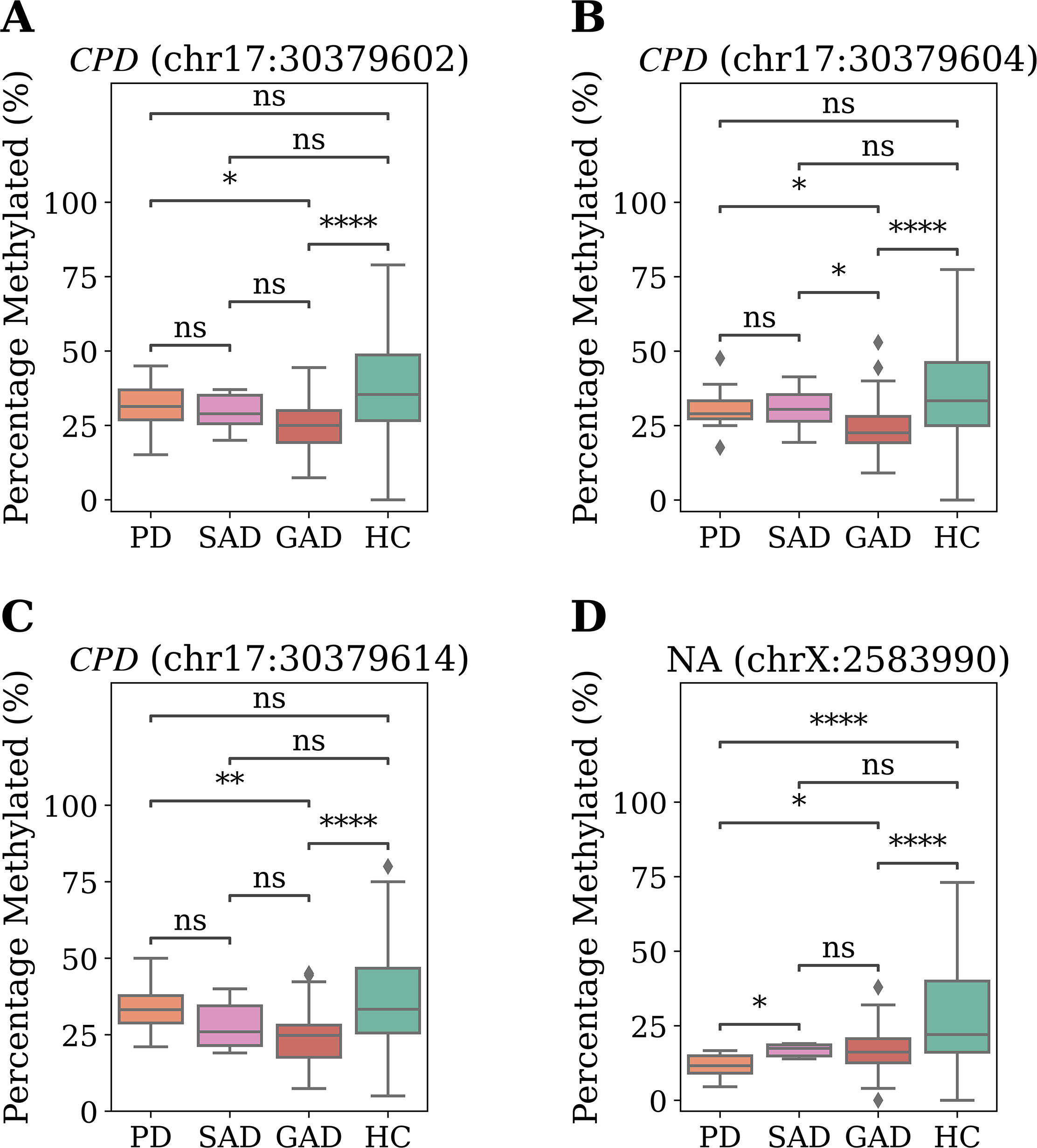
The biomarkers with significantly different methylation values across different types of anxiety disorders. The boxplot illustrates the methylation values of samples for four CpG sites that exhibited significant differences across different types of anxiety disorders. The position of each biomarker is indicated in the plot title, with significance levels denoted for differences between disease types. PD = Panic disorder; SAD = Social anxiety disorder; GAD = Generalized anxiety disorder; HC = Healthy control. A) chr17:30379602. B) chr17:30379604. C) chr17:30379614. D) chrX:2583990. ns: 5.00×10^-2^ < p ≤ 1.00×10^0^; *: 1.00×10^-2^ < p ≤ 5.00×10^-2^ ; **: 1.00×10^-3^ < p ≤ 1.00×10^-2^ ; ***: 1.00×10^-4^ < p ≤ 1.00×10^-3^ ; ****: p ≤ 1.00×10^-4^ ; Mann-Whitney-Wilcoxon test

In contrast, the biomarker on the X chromosome (chrX:2583990) indicated a statistically significant average methylation difference of 4.77% higher in PD patients in contrast to GAD patients (Fig. 3D, and Additional file 7: Table S5). It suggests that this hypomethylation pattern observed on chromosome X is present in GAD patients but appears more pronounced in patients with PD (Fig. 3D).

### Majority of 17 methylation biomarkers are located on CpG islands

Seventeen differentially methylated sites (DMSs) were located on the genomic locations of eleven genes (Table 2). Among these, 82.35% (14/17) of CpG sites were located in the promoter regions of genes (including 1∼5kb upstream of the transcription start site as promoter) (Fig. 4, and Table 2). Moreover, 82.35% (14/17) of CpG sites were located within CpG islands (Fig. 4). The other three CpG sites were also located on CpG shores (Fig. 4). All 17 methylation biomarkers identified in this study are located near CpG islands, as CpG shores are defined as 2kb upstream or downstream of CpG islands. Furthermore, all four regions where DMS biomarkers clustered within 100bp (*MIR3146*, *ZNF689*, *CPD*, and *MRPL45*) were located within CpG islands (Additional file 2: Fig. S3). This indicates that the biomarkers for anxiety disorders play a significant role in regulating molecular mechanisms within CpG-enriched regions.[33–35] In the pathway enrichment analysis, these eleven genes did not share any significant pathways (Additional file 8: Table S6).

**Table 2.** Summary of the 17 CpG site locations, associated genes, and CpG annotations.

| Chromosome | Position | Gene symbol | Genomic location | Direction of methylation difference | CpG Annotation |
| --- | --- | --- | --- | --- | --- |
| chr1 | 45339798 | *MUTYH* | Intron; Exon;<br>Far Promoter; 5'UTR;<br>Intron-Exon Boundary | Hypo | CpG Shore |
|  |  | *TOE1* | Promoter; Far Promoter |  |  |
| chr7 | 19708981 | *MIR3146* | Far Promoter | Hypo | CpG Island |
| chr7 | 19708988 | *MIR3146* | Far Promoter | Hypo | CpG Island |
| chr7 | 19708990 | *MIR3146* | Far Promoter | Hypo | CpG Island |
| chr10 | 632863 | *DIP2C* | Intron | Hypo | CpG Island |
| chr10 | 119892349 | *SEC23IP* | Promoter | Hypo | CpG Shore |
| chr10 | 132767837 | *INPP5A* | Intron | Hyper | CpG Shore |
| chr11 | 64304843 | *ESRRA* | Promoter; Far Promoter | Hypo | CpG Island |
| chr16 | 30610279 | *ZNF689* | Intron; Exon; Promoter;<br>5'UTR; Intron-Exon<br>Boundary | Hypo | CpG Island |
| chr16 | 30610288 | *ZNF689* | Intron; Exon; Promoter;<br>5'UTR; Intron-Exon<br>Boundary | Hypo | CpG Island |
| chr17 | 30379602 | *CPD* | Exon; Promoter | Hypo | CpG Island |
| chr17 | 30379604 | *CPD* | Exon; Promoter | Hypo | CpG Island |
| chr17 | 30379614 | *CPD* | Exon; Promoter | Hypo | CpG Island |
| chr17 | 38297134 | *MRPL45* | Exon; Promoter; 5'UTR | Hypo | CpG Island |
| chr17 | 38297137 | *MRPL45* | Exon; Promoter; 5'UTR | Hypo | CpG Island |
| chr18 | 46104880 | *ATP5F1A* | Promoter | Hypo | CpG Island |
| chrX | 2583990 | NA | NA | Hypo | CpG Island |

If a gene associated with a biomarker has annotations for different genomic locations based on transcripts, all genomic locations are listed with semicolons. The direction of methylation difference is indicated as "Hyper" for hypermethylated biomarkers and "Hypo" for hypomethylated biomarkers. NA = Not Available;

**Fig. 4.**
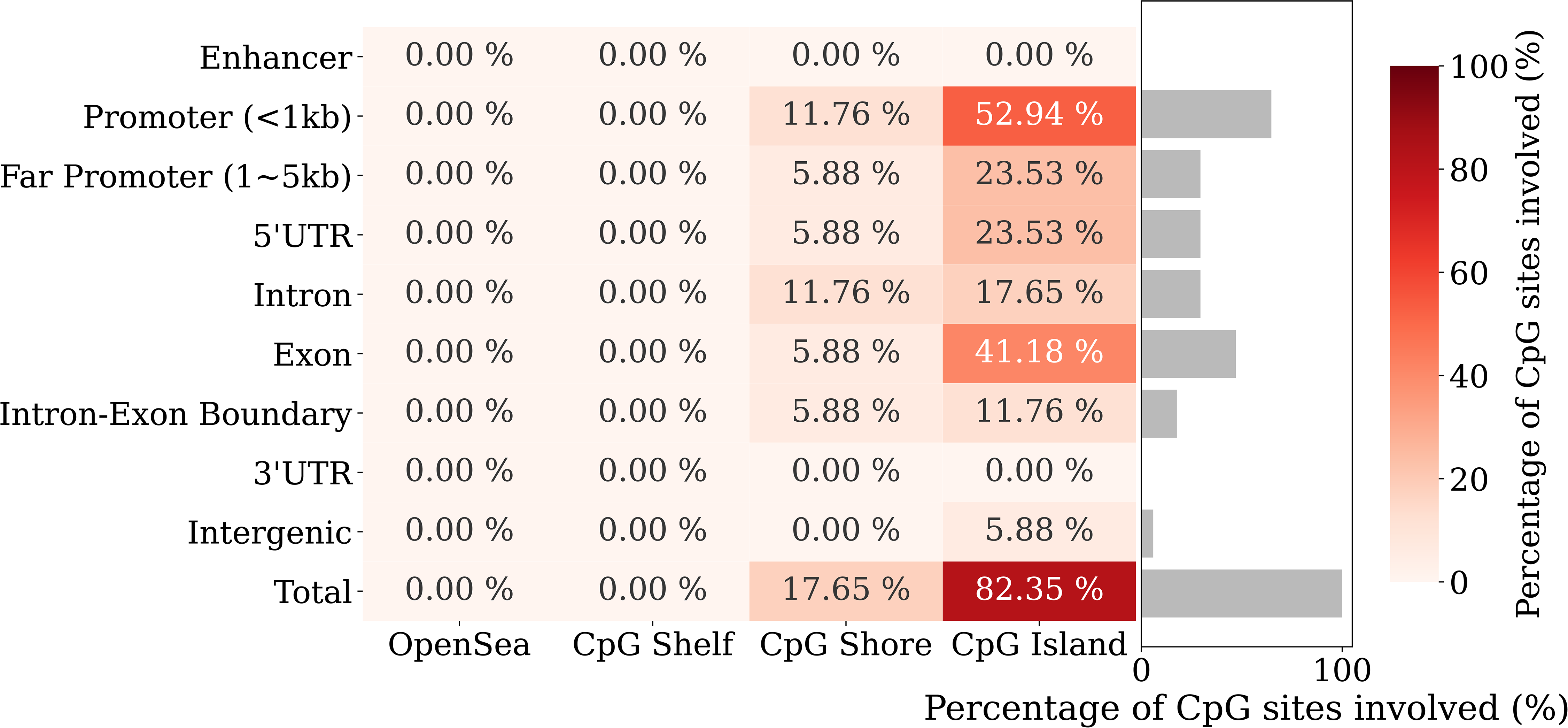
Summary of the genomic locations of the 17 biomarkers. Heatmap illustrates the frequencies of the 17 biomarkers based on their genomic location. The X-axis represents CpG annotation, while the Y-axis represents genomic annotation of genes. The heatmap colors, as indicated by the color bar on the right, depict the percentage involvement of 17 biomarkers across different CpG annotations and genomic locations. Bar plot shows the percentage of biomarkers in each genomic location.

### The interpretation of eleven genes associated with 17 methylation biomarkers

From the literature search, we found these genes are broadly associated with two main mechanisms: 1) mitochondrial dysfunction and cell apoptosis and 2) the regulation of neurosignaling. Five genes (*ATP5F1A*, *INPP5A*, *ESRRA*, *MRPL45*, and *MUTYH*) were related to mitochondrial functions, especially ATP synthesis and the response to Reactive Oxygen Species (ROS) stress (Fig. 5A, and Additional file 1: Supplementary Text).[36–43] Additionally, *ZNF689*, *MIR3146*, and *TOE1* were associated with cell apoptosis through cell stress (Fig. 5A, and Additional file 1: Supplementary Text).[44–49] In contrast, five genes (*MIR3146*, *ESRRA*, *SEC23IP*, *DIP2C*, and *CPD*(*SLC6A4*)) were directly related to serotonin and GABA neurotransmitters, which are known to be relevant to mood disorders like depression and anxiety (Fig. 5B, and Additional file 1: Supplementary Text).[44, 45, 50–62]

**Fig 5.**
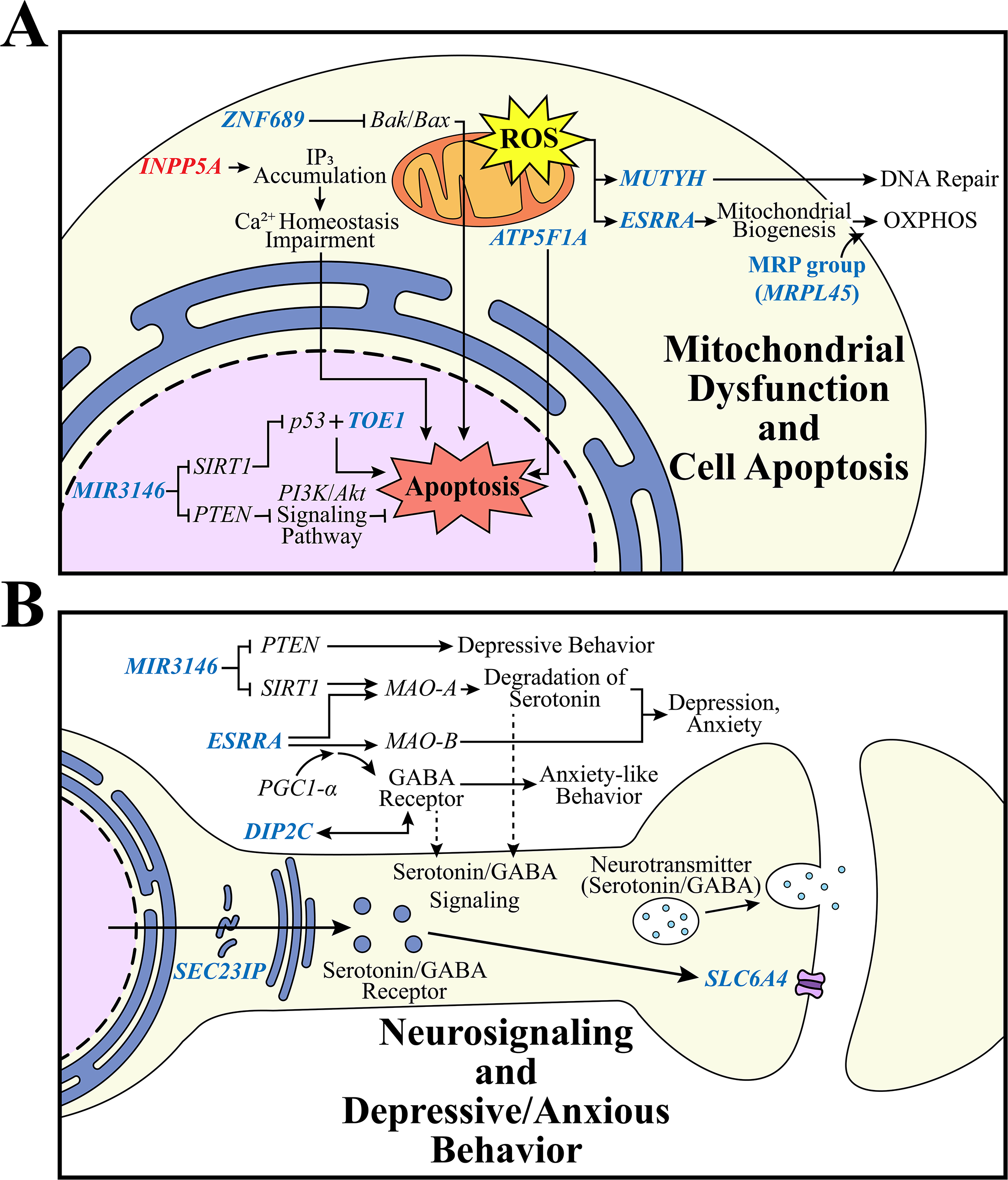
A diagram representing the molecular pathways of the 17 biomarkers’ associated genes. This figure represents the functions of genes in the cellular component and was drawn by the authors. The genes highlighed in blue represent genes identified as hypomethylation biomarkers in our study, while those in red denote genes identified as hypermethylation biomarkers. A) A chart illustrating the pathways of 8 genes related to mitochondrial dysfunction and cell apoptosis. The functions related to mitochondria and apoptosis are schematically illustrated, with major functions represented by star-shaped speech bubbles. ROS=Reactive Oxygen Species; OXPHOS=Oxidative Phosphorylation; B) A diagram depicting the pathways of 5 genes associated with serotonin/GABA neurotransmitter signaling and depressive/anxious behavior. The figure illustrates the serotonin/GABA-related pathway within nerve cells, depicting the transport of serotonin/GABA receptors from the endoplasmic reticulum exit site and the transmission of neurotransmitters.

It’s important to note that previous studies have demonstrated the impact of psychological stress on apoptosis in immune cells within the bloodstream.[63–68] This study, through the identification of differential methylation in genes associated with cell stress response, regulation of mitochondrial function, and control of apoptosis in individuals with anxiety disorders, provides support for these previous findings (Fig. 5A). The outcomes presented, along with the predominant localization of CpG biomarkers in CpG island regions observed in our investigation, indicate that the cell apoptosis pattern induced by anxiety disorders can instigate specific molecular alterations in the blood. We suggest this mechanism as a pivotal avenue for detecting anxiety disorders through blood-based assessments.

Furthermore, in this study, epigenetic changes in genes related to the regulation of neurotransmitters were observed in blood samples (Fig. 5B). The genes *MIR3146* and *ESRRA*, identified in this study, exhibit associations with both cell apoptosis and neurosignaling, implying a potential linkage between these two fundamental mechanisms.

Four out of eleven genes associated with methylation biomarkers have been reported as relevant in previous research.[10, 69–76] In previous studies, the altered methylation patterns of the *INPP5A* gene were associated with cognitive functions in both blood and the hippocampus samples.[69, 70] The *DIP2C* gene has been previously reported to undergo changes in its methylation patterns in studies related to anxiety disorder, post-traumatic stress disorder (PTSD), and major depressive disorder.[10, 71–74] Furthermore, despite their distinct genomic positions, both *INPP5A* and *DIP2C* have been identified as markers in a blood-based study for diagnosing anxiety disorders.[75] The methylation and gene expression patterns of the *ATP5F1A* gene are known to be associated with Alzheimer’s disease.[76] However, the majority of the genes associated with the remaining DMSs in this study have not previously been documented as undergoing methylation changes related to mental disorders.

The identified biomarker located in the intronic region of the *INPP5A* gene (chr10:132767837) in this study yields intriguing findings on multiple fronts. Noteworthy for being the sole hypermethylation biomarker among the 17 biomarkers and showcasing a remarkable 99.8% reproducibility, with only two exceptions in 1000 bootstrapping iterations, this biomarker has previously been linked to cognitive functions (Table 2, Additional file 6: Table S4).[69, 70] Furthermore, it ranked the highest importance in the XGBoost model (Fig. 2B). Collectively, these pieces of evidence underscore the potentially pivotal role of this gene in the landscape of anxiety disorders.

## Discussion

The anxiety patient classification model in our study overcame the limitations of relatively moderate performance observed in previous studies.[30–32] However, this study could not elucidate the specific causal relationships through which these biomarkers manifest uniquely in anxiety disorder patients. Additionally, all participants in this study already had anxiety disorders, making it impossible to determine whether these biomarkers had undergone changes prior to the onset of anxiety disorders. We think continuous monitoring of DNA methylation and clinical changes in individuals who did not have anxiety disorders before its onset can provide insights into the causality of the 17 biomarkers and reveal patterns of change associated with the progression of anxiety disorders.

As DNA methylation is known to control cell fate, it exhibits cell type-specific patterns.[77, 78] Therefore, the emergence of CpG sites related to neurosignaling as biomarkers in whole blood samples dominated by immune cells is distinctive. In a previous study, it has been established that the methylation pattern in blood provides limited information about the brain.[79] However, in our study, nine biomarkers were associated with mechanisms related to the regulation of neurotransmitter signaling pathways (Fig. 5B, Additional file 1: Supplementary Text). While two genes (*MIR3146* and *ESRRA*) concurrently involved in cell apoptosis and neurosignaling are thought to connect the two mechanisms, the reason for the emergence of such signals in the blood remains unexplained. Regardless, our study suggests the possibility that two mechanisms, mitochondrial dysfunction/cell apoptosis and neurosignaling, establish a causal relationship, playing a role in regulating each other.

The composition of the anxiety disorder patient sample group in our study is a significant limitation. The relatively small sample size of 94 patients may have led to a decrease in performance during the process of dividing the dataset into training and validation sets for biomarker discovery and validation. Furthermore, the fact that all patients were recruited from a specific hospital in Korea, with an average age of 42.98 years (Standard deviation: 15.64) (Table 1), suggests a high potential for biased results within a specific population. Given that DNA methylation can vary significantly based on ethnic group or age, recruiting a more diverse and larger sample group could provide a more general understanding of the characteristics of anxiety disorders.[80–82]

Finally, the biased distribution of patients across anxiety disorder types in our study, particularly favoring generalized anxiety disorder (GAD), is another limitation. As a result, the 17 biomarkers identified here reflect a bias towards GAD, and the statistically less significant differentiation of specific biomarkers based on other types of anxiety disorder. Although the biomarker located on chromosome X (chrX:2583990) was uniquely found to be more specific to panic disorder, the study lacks detailed explanations regarding why certain types of disorders exhibit more specific or less specific biomarkers (Fig. 3D). Understanding the reasons behind the emergence of more specific or less specific biomarkers in particular types of disorders is crucial, and additional analyses on this aspect would enhance the comprehensibility and interpretation of the study results.

## Conclusions

In our study, we identified and validated 17 novel epigenetic biomarkers associated with anxiety disorders, showing a performance of area under the curve (AUC) of 0.876. These biomarkers were linked to 1) mitochondrial dysfunction and cell apoptosis and 2) the regulation of neurosignaling. Despite the successful validation, this study could not explain the causality associated with the identified biomarkers in anxiety disorders, and it failed to interpret why biomarkers related to neurosignaling appeared in whole blood samples. Additionally, due to limitations in the anxiety patient dataset, the results in this study have the possibility of biased results toward a specific population. Further recruitment of additional sample cohorts is necessary to address these limitations. Nevertheless, despite these shortcomings, the study underscores the clinical utility of blood liquid biopsy in the diagnosis of anxiety disorders. Furthermore, these biomarkers indicate the potential for early diagnosis, predicting treatment efficacy, continuous monitoring, health screening, and delivering personalized therapeutic interventions for anxiety disorders.

## Declarations

### Ethics approval and consent to participate

Written informed consent was obtained from all participants in this study. Sample collection and sequencing were approved by the Institutional Review Board (IRB) of the Ulsan Medical Center (USH.20.013) and Ulsan National Institute of Science and Technology (UNISTIRB-15-19-A).

### Availability of data and materials

The datasets and materials used in the study are available from the corresponding author upon reasonable request.

## Author’s contributions

Conceptualization : S.J., B.K., H.J., E.S., and J.B.; Resources : J.A., H.U., Younghui K., H.B., and H.J.; Data curation : Yoonsung K., A.B., Y.J., Yeo Jin K., D.S., and H.J.; Methodology : Yoonsung K., A.B., Y.J., Yeo Jin K., K.A., S.J., H.R., D.S., E.S., and J.B.; Software, Formal analysis, Visualization, and Validation : Yoonsung K.; Investigation : Yoonsung K., A.B., Y.J., K.A., and D.S.; Funding acquisition : B.K., S.L., H.J., and J.B.; Supervision : S.L., H.J., E.S., and J.B.; Project administration : E.S., and J.B.; Writing-original draft : Yoonsung K., A.B., and D.S.; Writing-review&editing : Y.J., Yeo Jin K., K.A., S.J., H.R., S.L., H.J., E.S., and J.B.; All authors reviewed and approved the final version of the manuscript.

## List of abbreviations

AUC: Area under curve
AUPRC: Area under the precision-recall curve
AUROC: Area under the receiver operating characteristic curve
CpG site: A region of DNA where a cytosine is followed by a guanine that has the potential to be methylated
DASS-21: The Depression, Anxiety and Stress Scale - 21 Items DEGs : Differentially expressed genes
DMRs: Differentially methylated regions
DMSs: Differentially methylated sites EWAS : Epigenome-wide associdation study FDR : False discovery rate
GAD: Generalized anxiety disorder
GAD-2: Generalized Anxiety Disorder 2-item GO : Gene Ontology
HC: Healthy control Hyper : Hypermethylated Hypo : Hypomethylated
KEGG: Kyoto Encyclopedia of Genes and Genomes KGP : Korean Genome Project
MRS: Methylation risk score NA : Not available
OXPHOS: Oxidative Phosphorylation PD : Panic disorder
PRC: Precision-recall curve
PTSD: Post-traumatic stress disorder Reprod : Reproducibility
ROC: Receiver operating characteristic ROS : Reactive oxygen species
SAD: Social anxiety disorder
SHAP: SHapley Additive exPlanations TWAS : Transcriptome-wide association study

## Acknowledgments

We thank all the genome donating participants. The biospecimens for anxiety disorder patients were provided by the Ulsan Medical Center (USH.20.013). We thank the Korea Institute of Science and Technology Information (KISTI) which provided us with the Korean Research Environment Open NETwork (KREONET). We also appreciate the Ulsan ICT Promotion Agency (UIPA) which provided us with the BioDataFarm system which supports the storage, analysis, and management of the BioBigData.

## Supplementary Information

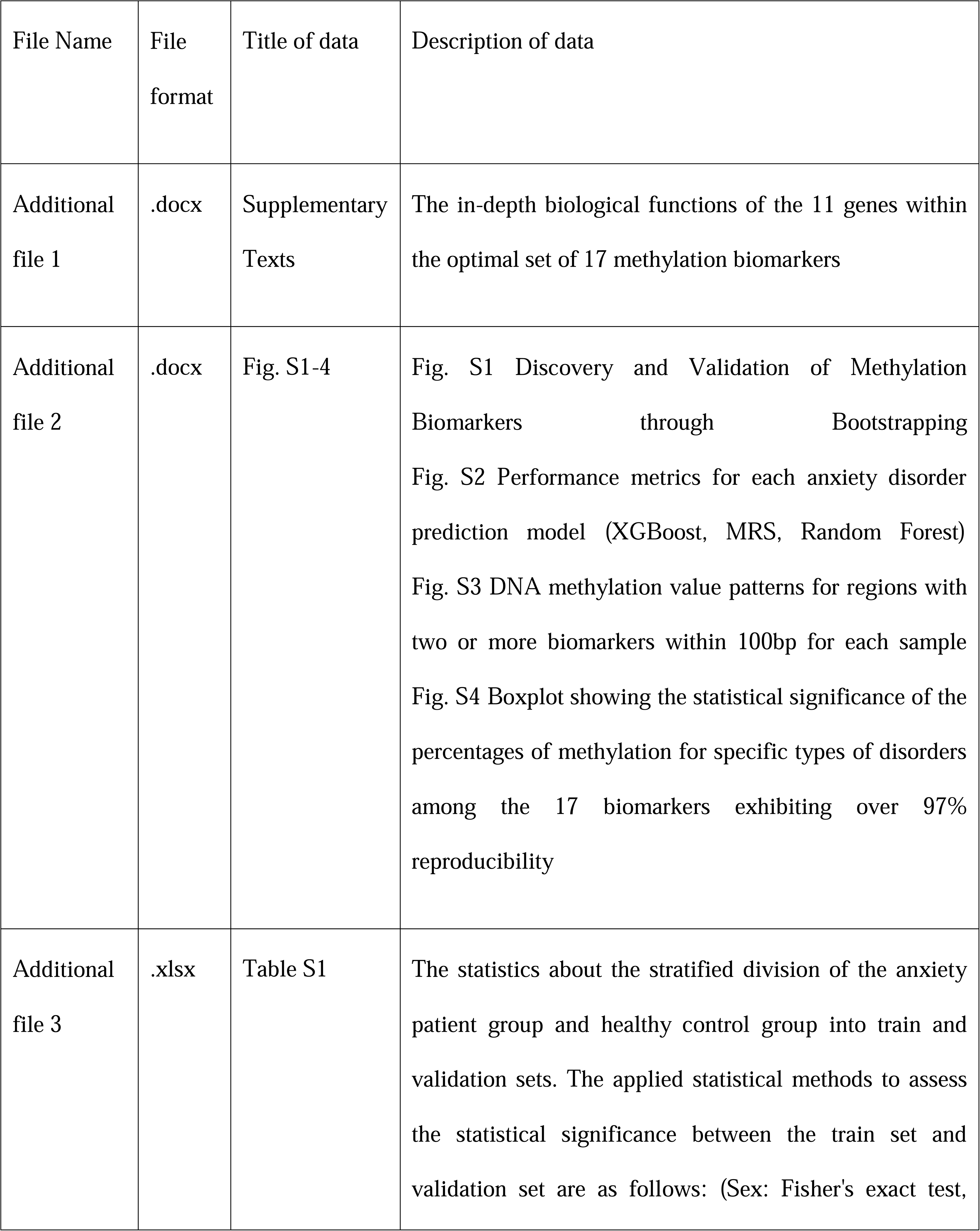
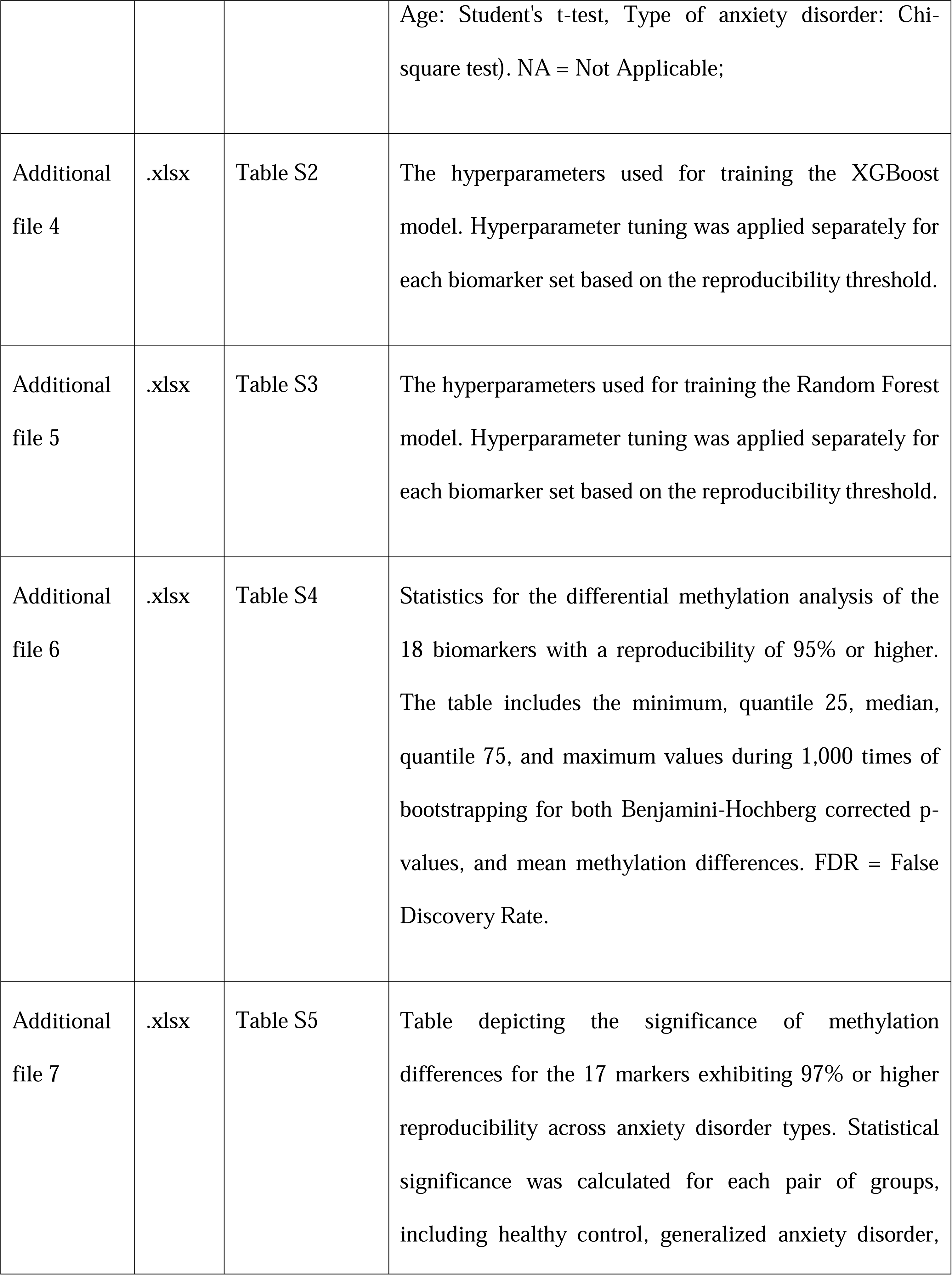
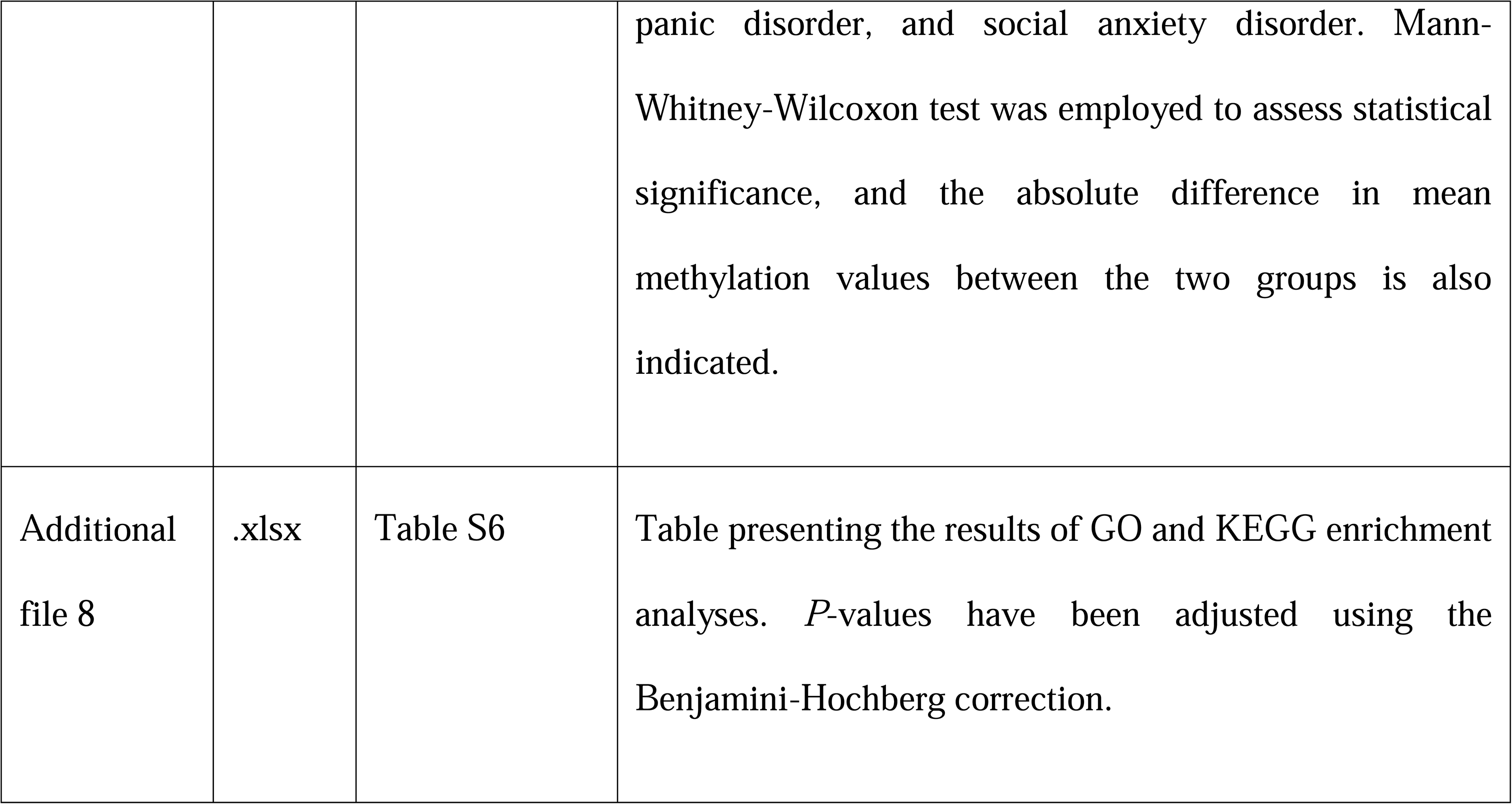

