## Supplementary Text for "Identification of 17 novel epigenetic biomarkers associated with anxiety disorders using differential methylation analysis followed by machine learning-based validation"

### **Supplementary Texts**

#### **The in-depth biological functions of the 11 genes within the optimal set of 17 methylation biomarkers**

In our identified biomarker set, five genes were associated with mitochondrial function. The ATP synthase (*ATP5F1A*), involved in energy production in mitochondria, was linked to the DMS biomarker.[1] Additionally, the *INPP5A* gene, which regulates calcium ion homeostasis through IP3 accumulation, was identified.[2-4] Moreover, ATP generation is closely linked to Reactive Oxygen Species (ROS) in mitochondria.[5] In our data, those genes related to ROS stress were associated with DMS biomarkers. *ESRRA* promotes mitochondrial biogenesis to reduce ROS stress[6], involving the MRP group (e.g., *MRPL45*) in Oxidative Phosphorylation (OXPHOS).[7] The *MUTYH* gene, known for repairing DNA damage induced by ROS stress, was related to the biomarker.[8, 9] Genes regulating cell apoptosis induced by ROS stress were also included.[10] *ZNF689*, well-studied in cancer research, regulates *Bcl-2* family genes, controlling apoptosis during mitochondrial stress.[11, 12] *MIR3146*, a microRNA, interacts with well-known apoptosis-regulating genes[13], *SIRT1*[14] and *PTEN*[15]. Additionally, the *TOE1* gene[16] stabilizes *p53* in the apoptotic signaling regulated by *SIRT1*.

Furthermore, five genes were directly associated with serotonin and GABA neurotransmitters, which are known to be relevant to mood disorders like depression and anxiety.[17-19] *MIR3146*, in conjunction with *SIRT1*[14] and *PTEN*[15], influences not only cell apoptosis but also regulation of the *MAO-A* gene.[20, 21] *ESRRA* was found to impact the expression of *MAO-A* and *MAO-B* genes[22, 23], which are known to be involved in neurotransmitter signaling and emotional regulation.[24-28] *SEC23IP* is linked to the transport of neurotransmitter receptors[29], while *DIP2C* is involved in the regulation of these receptors.[30] Notably, the DMS sites related to the *CPD* gene, positioned near *CPD*, *NF1*, and *SLC6A4*, have known associations in previous studies with these genes.[31] Among these, *SLC6A4* functions as a serotonin transporter, directly related to neurotransmitter signaling.[32]
