## Supplementary material for "Identification of 17 novel epigenetic biomarkers associated with anxiety disorders using differential methylation analysis followed by machine learning-based validation": Fig. S1, Fig.S2, Fig.S3, Fig.S4

### **Supplementary Figures**

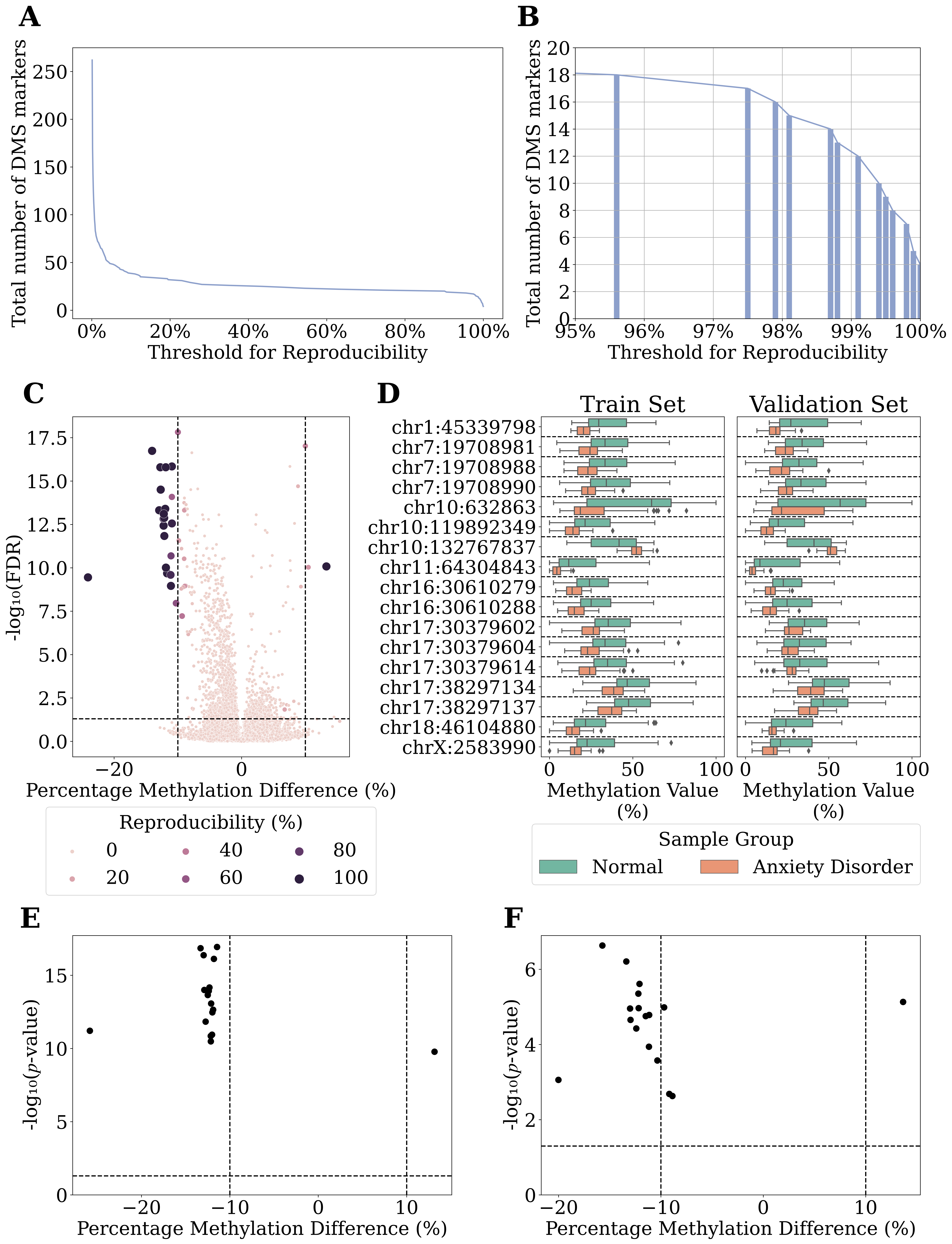

#### **Fig. S1 Discovery and Validation of Methylation Biomarkers through Bootstrapping.**

A) Line plot depicting the number of biomarkers based on the reproducibility threshold during 1000 rounds of bootstrapping. DMS = Differntially Methylated Sites; B) Cumulative plot illustrating the count of biomarkers showing 95% or higher reproducibility. Both the bar plot and line plot represent the number of biomarkers satisfying each threshold. DMS = Differntially Methylated Sites; C) Plot representing the reproducibility information for biomarkers in the training set during bootstrapping on the volcano plot displaying the overall Differentially Methylated Sites (DMS) calculated from the entire dataset. The darker the color and the larger the size of the dots, the higher the reproducibility. D) Boxplot showing the methylation values of 17 biomarkers demonstrating 97% or higher reproducibility in both healthy controls and anxiety disorder patients for both the train (left panel) and validation (right panel) sets. E) Plot highlighting only the 17 biomarkers with 97% or higher reproducibility in the volcano plot of the training set. F) Plot displaying only the 17 biomarkers with 97% or higher reproducibility in the volcano plot of the validation set.

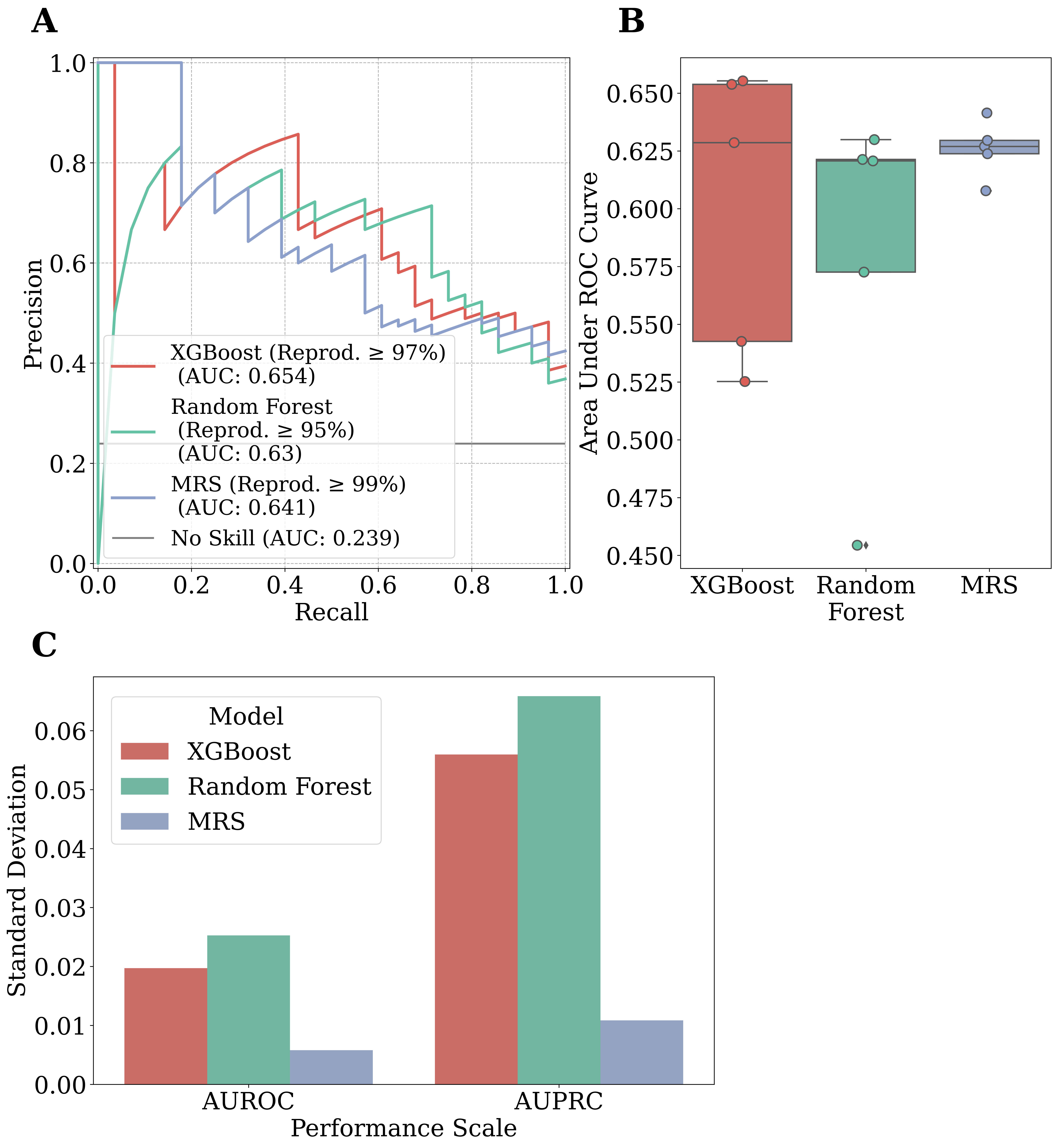

#### **Fig. S2 Performance metrics for each anxiety disorder prediction model (XGBoost, MRS, Random Forest).**

A) Precision-Recall (PR) curves for the biomarker sets with the highest Area Under the Receiver Operating Characteristic curve (AUROC) for each model type (XGBoost, Random Forest, MRS). Reprod = Reproducibility; B) Boxplot depicting the Area Under the PR Curve (AUPRC) values for each model type. Each dot represents the performance of the model for each of the five reproducibility thresholds. C) Barplot illustrating the standard deviation in performance (AUROC, AUPRC) across different biomarker sets for each model type (XGBoost , Random Forest, MRS).

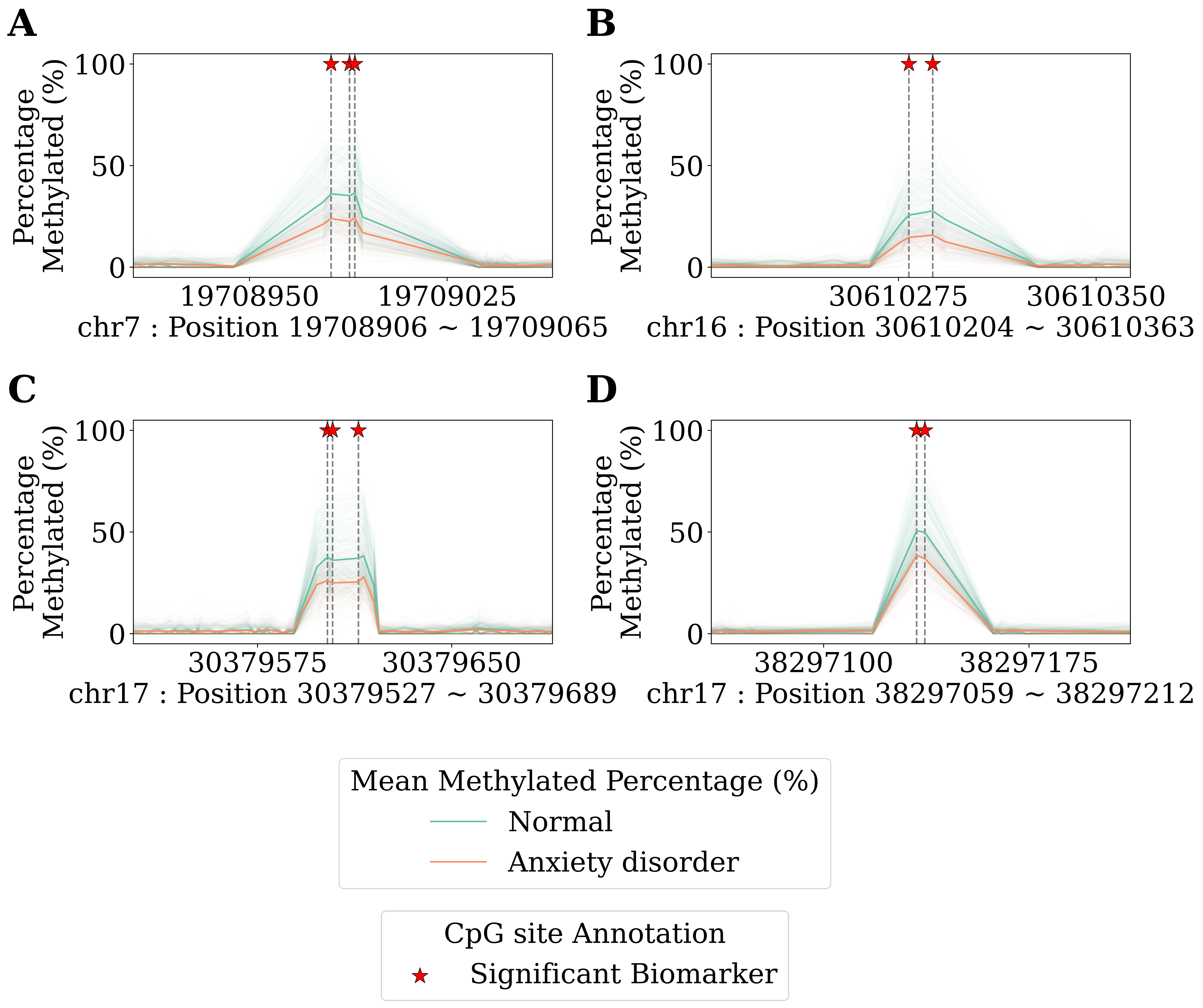

#### **Fig. S3 DNA methylation value patterns for regions with two or more biomarkers within 100bp for each sample.**

The green line represents healthy controls, the orange line represents anxiety disorder patients, with the bold line indicating the group's mean value and the faint line indicating values for each sample. The positions of designated CpG sites as biomarkers within the region are marked with star symbols. A) Chromosome 7: 19708881 ~ 19709090. B) Chromosome 16: 30610179 ~ 30610388. C) Chromosome 17: 30379502 ~ 30379714. D) Chromosome 17: 38297034 ~ 38297237

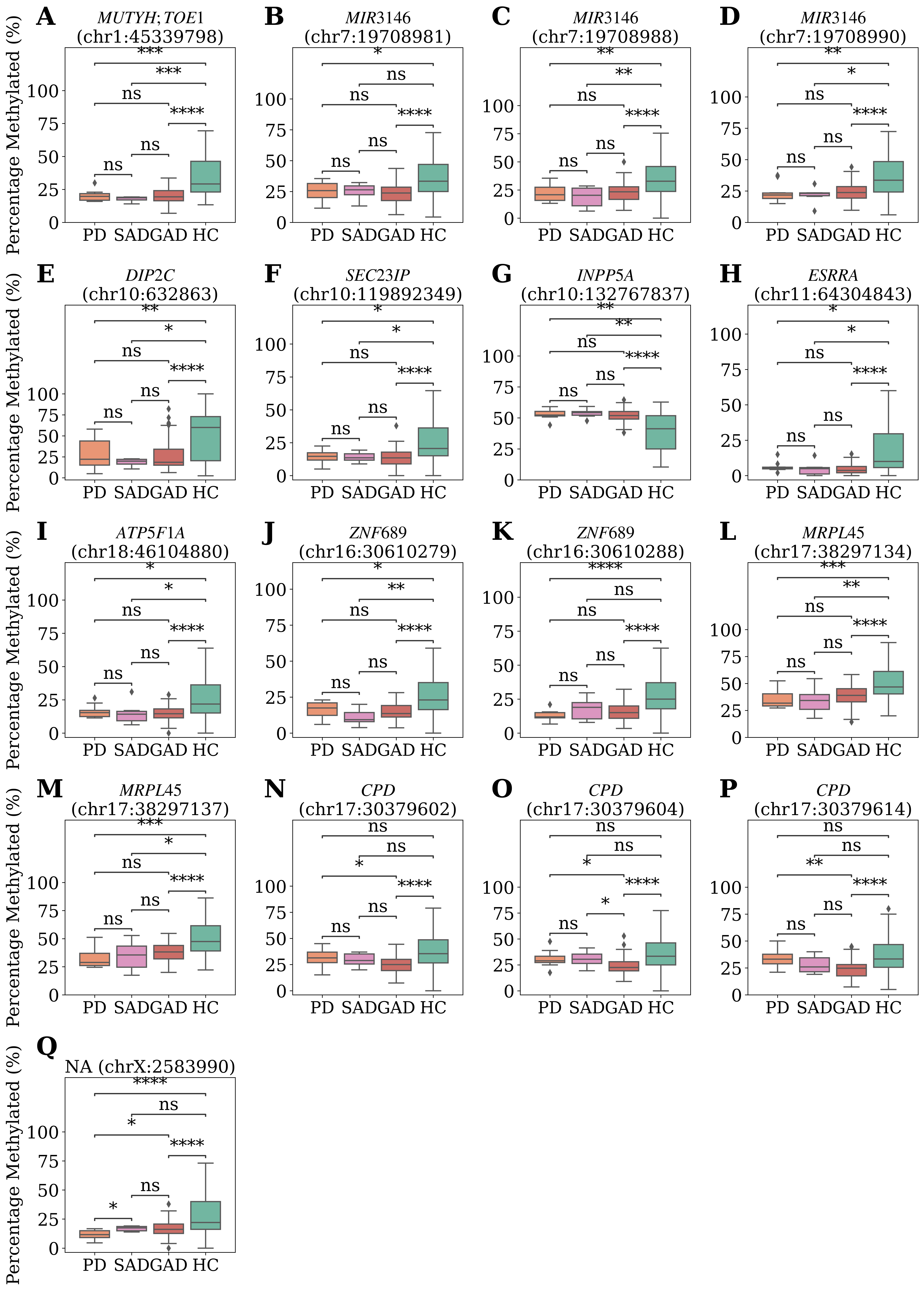

#### **Fig. S4 Boxplot showing the statistical significance of the percentages of methylation for specific types of disorders among the 17 biomarkers exhibiting over 97% reproducibility.**

The boxplot illustrates the methylation values of samples for each biomarker across different types of anxiety disorders. The position of each biomarker is indicated in the plot title, with significance levels denoted for differences between disease types. PD = Panic Disorder; SAD = Social Anxiety Disorder; GAD = Generalized Anxiety Disorder; HC = Healthy Control; A) chr1:45339798. B) chr7:19708981. C) chr7:19708988. D) chr7:19708990. E) chr10:632863. F) chr10:119892349. G) chr10:132767837. H) chr11:64304843. I) chr18:46104880. J) chr16:30610279. K) chr16:30610288. L) chr17:38297134. M) chr17:38297137. N) chr17:30379602. O) chr17:30379604. P) chr17:30379614. Q) chrX:2583990. ns: 5.00×10^-2^ < *p* ≤ 1.00×10^0^; *: 1.00×10^-2^ < *p* ≤ 5.00×10^-2^ ; **: 1.00×10^-3^ < *p* ≤ 1.00×10^-2^ ; ***: 1.00×10^-4^ < *p* ≤ 1.00×10^-3^ ; ****: *p* ≤ 1.00×10^-4^ ; Mann-Whitney-Wilcoxon test
